# Tunnel dynamics of quinone derivatives and its coupling to protein conformational rearrangements in respiratory complex I

**DOI:** 10.1101/2022.06.21.497056

**Authors:** Jonathan Lasham, Outi Haapanen, Volker Zickermann, Vivek Sharma

**Affiliations:** Department of Physics, University of Helsinki, 00014 Helsinki, Finland; Institute of Biochemistry II, University Hospital, Goethe University, 60438 Frankfurt am Main, Germany; Centre for Biomolecular Magnetic Resonance, Institute for Biophysical Chemistry, Goethe University, 60438 Frankfurt am Main, Germany; HiLIFE Institute of Biotechnology, University of Helsinki, 00014 Helsinki, Finland

## Abstract

Respiratory complex I in mitochondria and bacteria catalyzes the transfer of electrons from NADH to quinone (Q). The free energy available from the reaction is used to pump protons and to establish a membrane proton electrochemical gradient, which drives ATP synthesis. Even though several high-resolution structures of complex I have been resolved, how Q reduction is linked with proton pumping, remains unknown. Here, microsecond long molecular dynamics (MD) simulations were performed on *Yarrowia lipolytica* complex I structures where Q molecules have been resolved in the ~30 Å long Q tunnel. MD simulations of several different redox/protonation states of Q reveal the coupling between the Q dynamics and the restructuring of conserved loops and ion pairs. Oxidized quinone stabilizes towards the N2 FeS cluster, a binding mode not previously described in *Yarrowia lipolytica* complex I structures. On the other hand, reduced (and protonated) species tend to diffuse towards the Q binding sites closer to the tunnel entrance. Mechanistic and physiological relevance of these results are discussed.

## Introduction

Respiratory complex I is the first electron acceptor in many bacterial and mitochondrial electron transport chains, and its catalytic mechanism involves the reduction of quinone (Q) from NADH. The energy gain from Q reduction is used to pump protons across the inner mitochondrial membrane leading to the formation of an electrochemical gradient (Fig. 1A), which powers ATP generation (Agip, Blaza, Fedor, & Hirst, 2019; Kaila, 2018; Sazanov, 2015; Yoga, Angerer, Parey, & Zickermann, 2020). How exactly the reactions at the active site of complex I are coupled to proton pumping some 200 Å away remains a mystery. Computational studies have indicated the role of electrostatics and conformational dynamics, protein hydration and Q binding in the long-range electron-proton coupling in complex I (Galemou Yoga, Schiller, & Zickermann, 2021; Haapanen, Reidelbach, & Sharma, 2020; Haapanen & Sharma, 2021). Recent high resolution structural data from cryo electron microscopy (Chung et al., 2022; Grba & Hirst, 2020; Gu, Liu, Guo, Zhang, & Yang, 2022; Kampjut & Sazanov, 2020; Parey et al., 2018; Parey et al., 2019; Parey et al., 2021; Yoga, Parey, et al., 2020) have provided new insights into the role of Q binding, loop dynamics and water molecules in proton pumping by complex I.

**Fig. 1:**
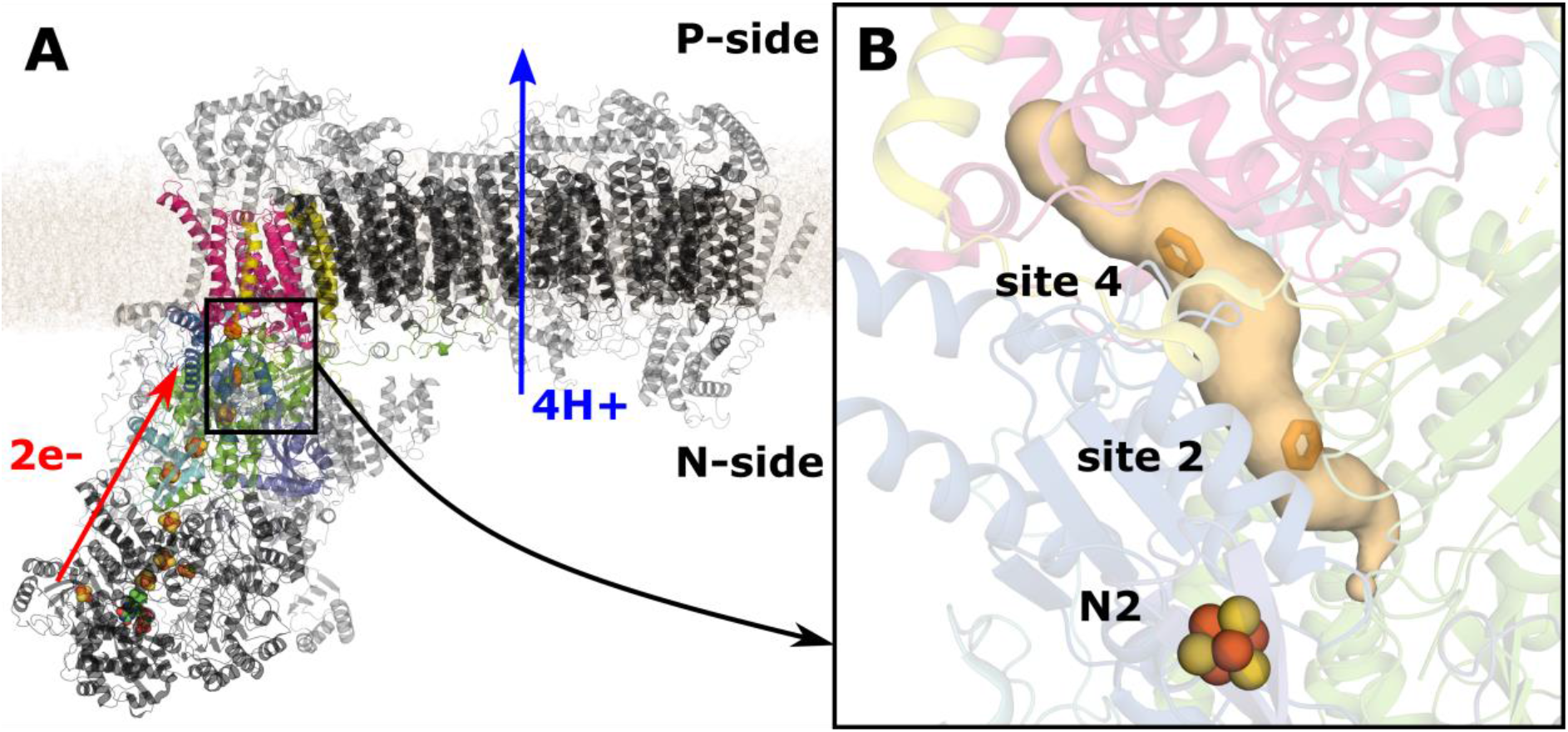
Respiratory complex I and its Q binding sites. **A** shows the entire structure of complex I from *Y. lipolytica* (PDB 6RFR) embedded in a lipid bilayer. Subunits close to the Q binding domain are shown in colors. NDUFS3 is shown in dark blue, NDUFS2 in green, NDUFS8 in cyan, NDUFS7 in blue, ND1 in magenta, and ND3 in yellow. Iron sulfur clusters buried in protein matrix are shown as orange and yellow spheres, and FMN is shown as green spheres. The inset **B** shows the Q binding tunnel as a light orange surface, with the Q head at sites 2 and 4 shown as orange licorice. The N2 FeS cluster is shown. The site 2 and site 4 positions of Q are based on PDB 7O6Y and PDB 6RFR, respectively. The tunnel was calculated using software CAVER (Pavelka et al., 2012) from PDB 7O6Y with a probe radius of 0.6 Å.

The Q molecule consists of a polar aromatic head and a long hydrophobic tail, and it binds in a ~30 Å long cavity known as the Q tunnel in complex I. The head, which undergoes redox reactions at the reaction site near the N2 FeS cluster (Fig. 1), can exist in several different redox and protonation states. The tail consists of multiple isoprene units of varying lengths depending on the species and helps in anchoring and guiding the Q within the long tunnel (Fedor, Jones, Di Luca, Kaila, & Hirst, 2017). Based on umbrella sampling and unbiased MD simulations, five distinct Q binding sites were proposed (Haapanen, Djurabekova, & Sharma, 2019; Teixeira & Arantes, 2019; Warnau et al., 2018). Out of the five sites, two were identified at the interface of the membrane and the peripheral arms of complex I (called sites 4 and 5). Latest high-resolution cryo EM data confirmed the existence of these sites (Kampjut & Sazanov, 2020; Parey et al., 2019). However, their functional meaning remains unclear, either they represent transient halts for Q upon its travel to and from the active site near N2 FeS cluster or they have a role in coupling Q-tunnel redox reactions to proton pumping in the membrane arm of complex I (Djurabekova et al., 2022; Haapanen & Sharma, 2021; Wikstrom, Sharma, Kaila, Hosler, & Hummer, 2015).

In addition, the two Q binding sites (1 and 2) closer to the N2 FeS center are found at the interface of NDUFS2 and NDUFS7 subunits. At these sites Q is expected to be reduced by electron transfer(s) from N2. Both sites have been confirmed by structural data (Chung et al., 2022; Gu et al., 2022; Gutiérrez-Fernández et al., 2020; Kampjut & Sazanov, 2020; Parey et al., 2021) as well as MD simulations (Haapanen et al., 2019; Warnau et al., 2018).

At the Q binding site 1, Q head group makes a hydrogen bond to Tyr144 of NDUFS2 subunit, which is known to be functionally important for Q redox reactions from mutagenesis studies (Tocilescu et al., 2010). Computational work suggests redox-coupled proton transfer reaction of Q bound at site 1 converts it to QH2 (or anionic QH-)(Sharma et al., 2015), which diffuses to site 2 upon conformational changes in the site, in particular in the β1-β2 loop of NDUFS2 subunit (Haapanen et al., 2019; Tocilescu et al., 2010; Warnau et al., 2018). The site 2 corresponds to a position where a Q molecule is not making a direct hydrogen bond to Tyr144.

The journey between these tunnel-bound Q sites, and in and out of the Q tunnel, is thought to be dependent on Q-tail length (Fedor et al., 2017; Haapanen et al., 2019), changes in protein environment, including sidechain movements (Haapanen et al., 2019; Yoga et al., 2019) as well as on changes in tunnel hydration (Teixeira & Arantes, 2019). However, how exactly these different aspects drive dynamics of different Q species in practice remains unclear. In particular, the role of protein-Q interactions and protein-protein interactions is poorly understood.

In the present work, we use long time-scale atomistic MD simulations on *Y. lipolytica* complex I, where Q has been structurally resolved at sites 2 and 4 (Fig. 1B), to investigate how different redox states of Q behave in the Q tunnel, and how this is coupled to changes in the protein conformation. Three different cryo-EM structures from *Y. lipolytica* were simulated (see methods): PDB 6RFR (Parey et al., 2019), which has a Q resolved at site 4 (setup S1), PDB 6GCS (Parey et al., 2018) with Q modeled at site 2 (setup S2), and finally PDB 7O6Y (Parey et al., 2021) which also has Q at site 2, although positioned slightly closer to the N2 cluster (setup S3). Each of these structures was simulated with four different states of Q: fully oxidized quinone (Qox), anionic semiquinone (SQ-), neutral semiquinone (SQ), and reduced and doubly protonated quinol (QH2). The simulations show dependence of Q state on its binding within in the Q tunnel, and that its diffusion between tunnel-bound sites is coupled to both loop dynamics and the formation and dissociation events of conserved ion pairs.

## Results

### Dynamics of Q in its different redox and protonation states

Figure 2 shows the distance of the Q head group from the N2 cluster during different simulations, with the starting positions marked by a pink dotted line. The simulations from setup S1, where Q was modeled at site 4, show a lot of similarity between the different Q species. However, it is notable that Qox shows two stable positions at 28 and 31 Å from N2 cluster, while QH2 only shows one stable position at ~27 Å, remarkably close to the structural position (PDB 6RFR). This raises the possibility that in structure a higher fraction of Q observed at site 4 may be the reduced and protonated quinol. The two radical SQ species (anionic and neutral) both show overall similar binding distances to Qox, however there are some notable instances where anionic SQ moves briefly towards the N2 cluster (Fig. S1).

**Fig. 2:**
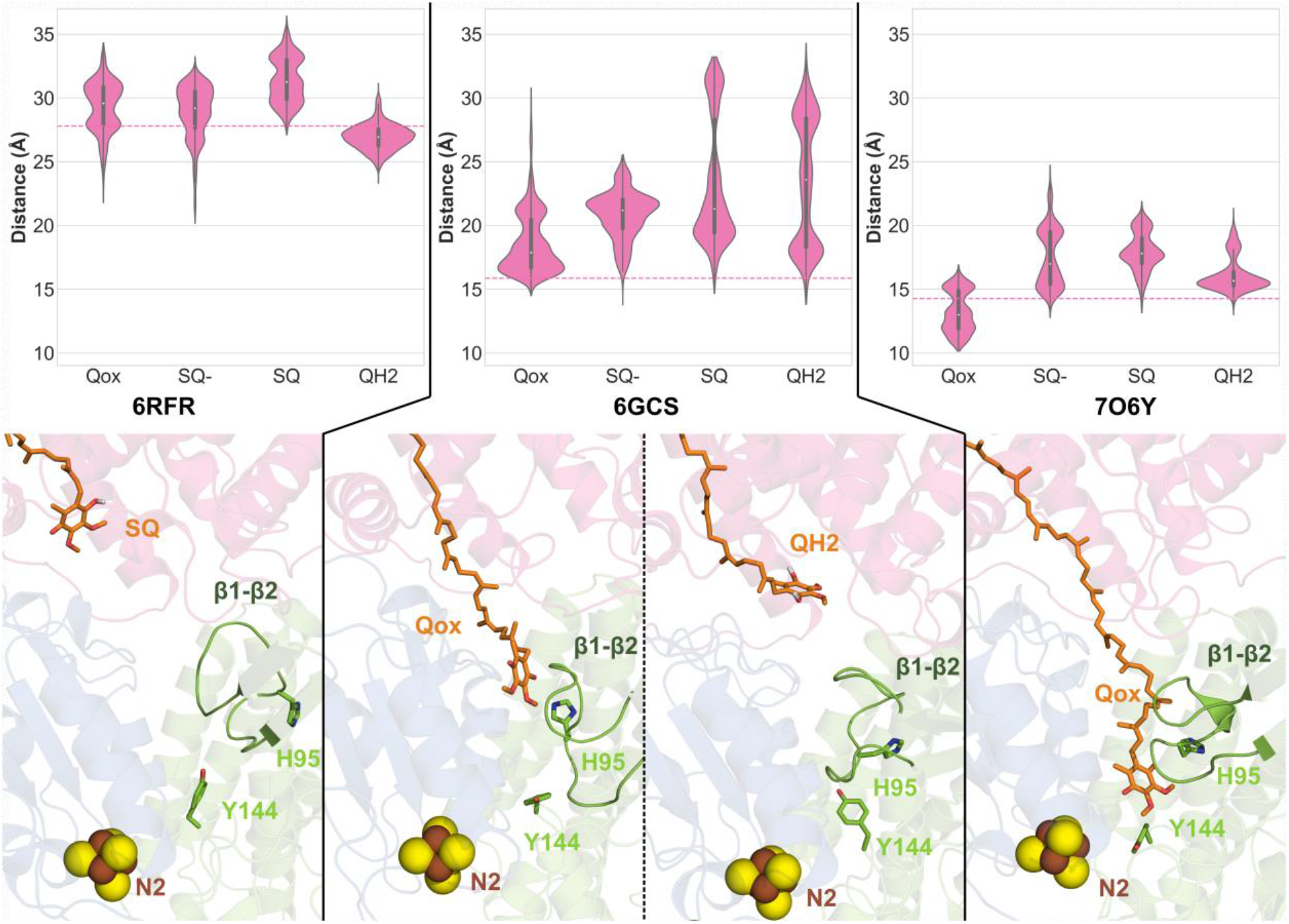
Violin plots showing the distance between the Q head group and N2 cluster for four different Q species. The three plots show simulation data from three separate structures: 6RFR (S1) where Q is modeled at site 4, 6GCS (S2) where Q is modeled at site 2, and 7O6Y (S3) where Q is also modelled at site 2. The pink dotted line represents the position of the Q head group observed in the structures. The lower panels show snapshots from various simulations. The Q molecule is shown in licorice, and the β1-β2^NDUFS2^ loop position is highlighted. Key conserved residues associated with Q binding, Y144^NDUFS2^ and H95^NDUFS2^, are shown in licorice.

Conversely, the simulations of setup 2 (PDB 6GCS) with Q modeled at site 2 show a much clearer dependence of redox state on Q-N2 distance during simulations than setup 1. Here, Qox is quite stable at site 2, with a major population close to the starting position, whereas QH2 is much more dynamic. In 2 out of 3 simulation replicas, QH2 moved away from site 2 and stabilized close to site 4 (Fig. S1). The Q-N2 distance at site 4 measured in these simulations is around 28 Å, which is remarkably close to the stable position from the setup S1 simulations. It is to emphasize that this is also in agreement with earlier estimates from umbrella sampling simulations of QH2 being stable (more than Qox) at site 4 of the Q tunnel (Warnau et al., 2018).

The subsequent site 2 simulations using the higher-resolution structure, PDB 7O6Y (setup S3), reveal differing behavior for Qox and QH2 compared to the setup S2 simulations. Overall, all Q species show higher stability at site 2, and there are no instances of Q moving towards site 4. In 2 out of 3 replicas, however, Qox moves closer to the N2 cluster towards site 1 (Fig. 2). This position of Q has previously been observed in bacterial and mammalian complex I structures (Chung et al., 2022; Gu et al., 2022; Gutiérrez-Fernández et al., 2020), but not in *Y. lipolytica* complex I structural data. Here, our MD simulations show that oxidized Q (Qox) can indeed bind closer to the N2 FeS cluster also in *Yarrowia* complex I, which may enhance efficiency of electron transfer from N2 to Q (Moser, Farid, Chobot, & Dutton, 2006).

In the S2 simulations, the radical semiquinone species (anionic) modelled at site 2 tend to move from the starting position of 16 Å to a position around 20 Å from the N2 cluster. Interestingly, the neutral SQ species diffuses even further towards the entrance of the Q tunnel and shows stable binding at ~31 Å, closer to the Q binding sites 4 and 5. This indicates that the neutral SQ species is much more mobile in the Q tunnel compared to anionic SQ-. Simulations on the higher resolution structure, PDB 7O6Y (setup S3), also show that the radical SQ species shifts slightly from the site 2 position to around 20 Å distance from N2. However, neutral SQ did not move further from this position towards site 4, reflecting relative stability of neutral SQ (and also QH2) in S3 simulations.

We next analyzed the possible source of this differing behavior of the Q species in the two different MD setups S2 and S3. We find that the position of the conserved β1-β2^NDUFS2^ loop (Figure 2, opaque green loop) is central factor deciding for Q dynamics in two setups (Figure S2). In the S2 runs based on PDB 6GCS, the loop is modelled (see Materials and methods) with His95 positioned in front of the Q head, meaning access to site 1 is blocked. On the other hand, in PDB 7O6Y, the loop is resolved with His95 pointing to the side of the Q headgroup, which means Q can more readily access site 1, as seen in the Qox state simulations. In addition, His95 blocking site 1 triggers Q movement away from site 2, as seen in the SQ and QH2 state simulations in setup S2. This also explains the lack of movement of neutral SQ and QH2 from site 2 to site 4 in the S3 simulations. An analysis of all S2 and S3 simulations show higher fluctuations of the β1-β2 loop to be coupled with Q movement (Figure S2). Overall, our data indicate that the movement of Q species is tightly coupled to β1-β2 loop position and dynamics.

### Interactions between quinone and protein (Q-protein interactions)

The heatmap in Fig. 3 shows percentage of the simulation time that different residues were in contact with the different Q states from the S2 simulations. Only three residues show consistently strong Q-protein interactions with each of the redox/protonation state studied (Met195^NDUFS2^, Phe203^NDUFS2^ and Met91^NDUFS7^). We point out that hydrophobic Met91^NDUFS7^ is well-known to be a residue central for Q binding and dynamics (Angerer et al., 2012; Fendel, Tocilescu, Kerscher, & Brandt, 2008; Haapanen et al., 2019; Parey et al., 2021). Interestingly, Qox retains the most contacts with the residues that were in contact at the beginning of the simulation, while the other states make newer and more transient interactions. This reflects the Q-N2 distances from Fig 2, which showed Qox to be most stable at site 2 in S2 simulations based on PDB 6GCS. The stability of Qox is partly explained by a hydrogen bond between the Q head group and His95 from the β1-β2 loop, which was observed for 28% of the total simulation time. Interactions in the S3 simulations were similar to this in all redox states, however hydrogen bonds to H95 were not observed due to its different orientation in the structure. (Figure S3B).

**Fig. 3:**
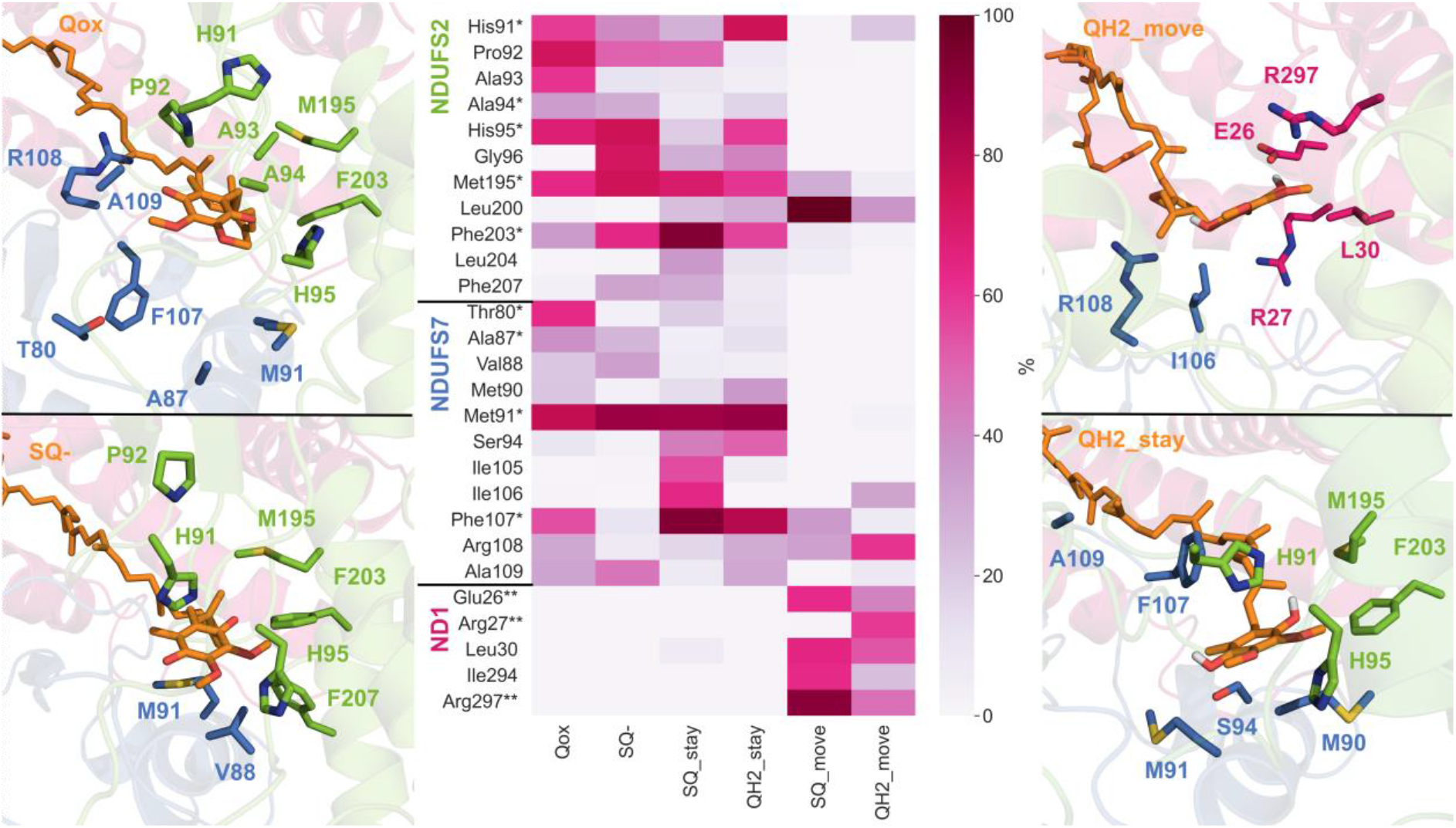
Interactions of protein residues within 5 Å of Q head group in S2 MD simulations. The color gradient from white to dark purple indicates the percentage of the trajectory data where the interaction is present. The heatmap is shown for Q redox/protonation states studied in this work. Here, ‘stay’ refers to selected frames of the trajectory where Q head group is less than 25 Å from N2, while ‘move’ refers to frames where the distance is more than 25 Å. A single asterisk (*) represents interactions present in the structure with Q resolved at site 2, while a double asterisk (**) indicates an interaction present in the structure with Q at site 4.

The Q-protein interactions of anionic SQ-in S2 simulations are partly similar to Qox, however SQ- makes additional contacts with Gly96^NDUFS2^ and Phe207^NDUFS2^, as well as Val88^NDUFS7^, indicating it binds in a slightly different way to Qox, which is reflected in the different Q-N2 distances (Fig. 2). Interestingly, both residues Phe207^NDUFS2^ and Val88^NDUFS7^ upon mutation are known to affect complex I activity (Angerer et al., 2012; Fendel et al., 2008). Moreover, many of the original interactions are maintained in the SQ-simulations, even when Q moves from its original position, resulting from many interacting residues being located on flexible loops facing Q tunnel.

Since QH2 and neutral SQ are most mobile in the S2 simulations, the contact analysis was broken down into two groups: when the Q-N2 distance is less than 25 Å, and when the Q-N2 distance is more than 25 Å. This roughly corresponds to Q staying at site 2 and Q leaving site 2 towards site 4 towards the tunnel entrance, respectively. When QH2 stays at site 2, the Q-protein interactions overall resemble Qox, with many of the interactions that were present in the beginning being stable. In addition, stable hydrogen bonds are seen between QH2 and His91^NDUFS2^ and His95^NDUFS2^ of the β1-β2^NDUFS2^ loop, when it stays at site 2. In contrast, neutral SQ shows some clear differences, and it has a relatively weak interaction to His91 and His95 of the β1-β2 loop based on contact analysis (Fig. 3). Neutral SQ’s inability to make stable interactions to these catalytically important histidine residues may be the reason for its instability at site 2 and explain its movement away from the structural binding position. In contrast, the anionic SQ-species is seen to anchor to site 2 by forming a stable hydrogen bond with His95 (ca. 35 %).

When both neutral SQ and QH2 move towards site 4, new contacts are established with protein residues from the membrane-bound ND1 subunit. Many of these contacts are also seen in the structurally resolved site 4 position from PDB 6RFR (denoted by double asterisk ** in the heatmap in Fig. 3). Significantly, some hydrophobic residues from the NDUFS7 loop (Ile106^NDUFS7^, Phe107^NDUFS7^) interact with different Q species at both site 2 and site 4, which indicates they may be of functional relevance. These two residues have indeed been identified in prior biochemical and computational studies to be important (Yoga et al., 2019). Another key residue which shows interactions at both site 2 and site 4 is Arg108^NDUFS7^. In Qox simulations, these interactions occur when Q is close to site 2, however SQ and QH2 do not form a stable interaction with Arg108^NDUFS7^ until they are moving towards site 4. The interactions between Q and Arg108^NDUFS7^ have been observed in both structures and simulations at site 4/5 (Haapanen et al., 2019; Kampjut & Sazanov, 2020; Parey et al., 2019), and mutation of arginine to glutamate is known to stall Q dynamics in the Q tunnel (Yoga et al., 2019). In addition, interactions with conserved Leu200^NDUFS2^ are also present when SQ and QH2 move towards site 4, which suggests this residue may also play an important functional role.

The interactions between Q and protein are also quite stable in the simulations with Q modeled at site 4 (setup S1, Figure S3A). Stable interactions with Ile106^NDUFS7^ and Arg27^ND1^ are seen with all four of the quinone species. However, many unique interactions are also present for each of the species, and this mirrors the difference seen in Q-N2 distances. Qox, SQ, and SQ-are all able to make stable interactions to Arg108^NDUFS7^, Phe224 ^ND1^ and Phe228^ND1^. Although not explicitly modelled in MD, transient π–π stacking-like interactions are also seen in the simulations between the head group of Qox and Phe228^ND1^. In a recent study, similar stacking interactions have been reported to form between the conserved Phe228^ND1^ residue and benzene ring of artificial quinone compounds, highlighting the importance of aromatic residues in trapping Q in the tunnel (Uno et al., 2022). In addition, SQ and SQ- make additional interactions to Trp77^NDUFS7^ and Leu57^ND1^. Interestingly, SQ is the only species found to interact with Tyr232^ND1^, by forming a stable hydrogen bond with Tyr232^ND1^ via Thr23^ND1^. Overall, the interaction analysis presented here highlights the role of several amino acid residues that interacts with Q upon its binding and dynamics in the Q tunnel (Table S1).

### Ion-pair dynamics coupled to Q movement

In addition to protein-Q interactions discussed above, several protein-protein interactions were also identified, which appear to depend on the binding position of the Q molecule. Trajectory data was analyzed from the S2 simulations, and data from Qox and QH2 simulations where Q was stable at site 2 were compared to data from QH2 simulations where Q moved towards site 4. Fig. 4 shows the sidechain distance of various ion pairs for these three data sets. The plots indicate that there is preference for certain ion pairs when Q stays at site 2 (left orange panels), with the other ion pairs preferentially forming when Q migrates towards site 4 (right cyan panels). Snapshots representative of the two situations are also shown.

**Fig. 4:**
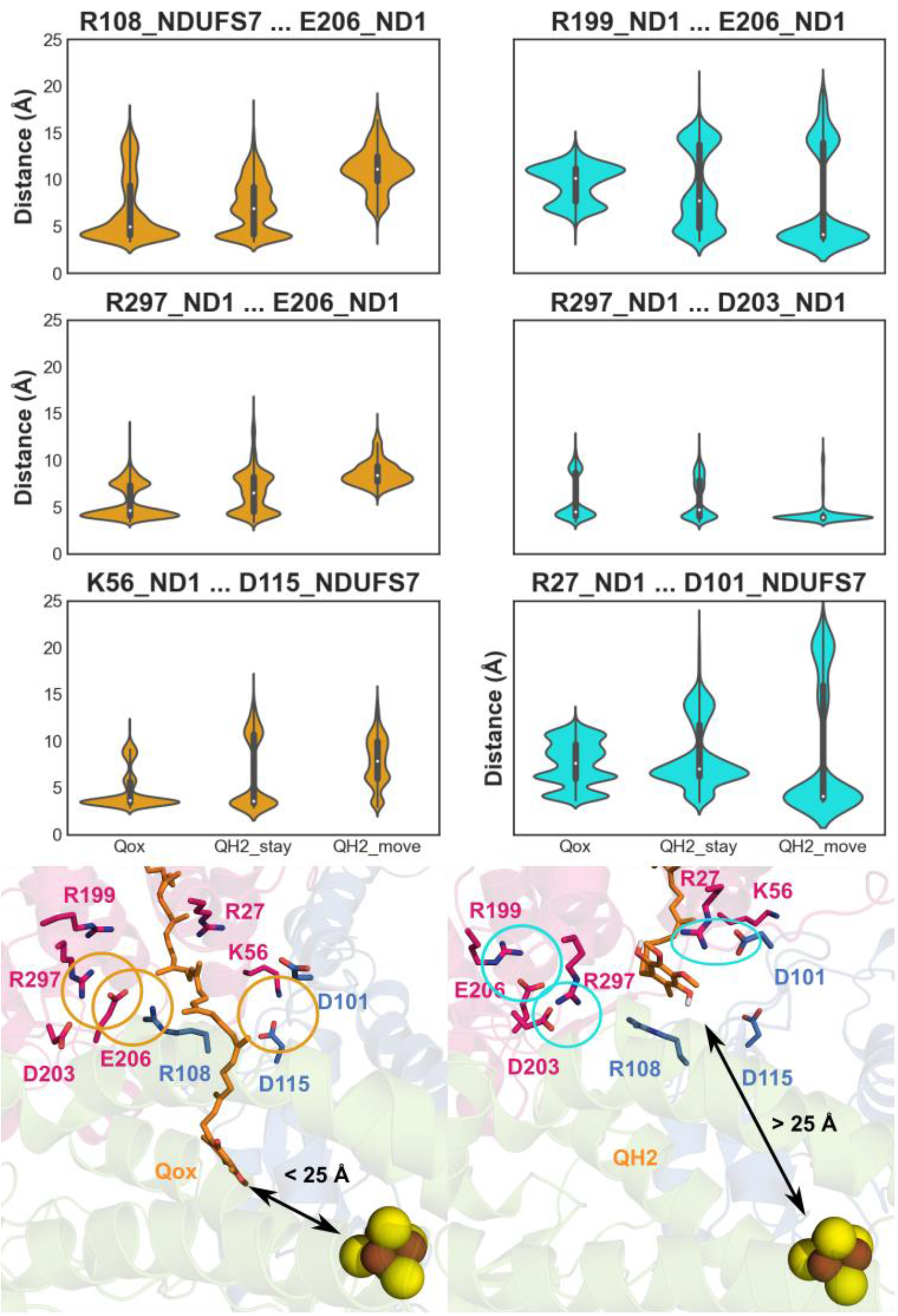
Sidechain distance of various ion pairs throughout trajectories shown as violin plots. Qox refers to site 2 simulations where oxidized Q (Qox) was modeled and simulated (setup S2). QH2_stay refers to frames from site 2-based MD simulations where the Q-N2 distance was less than 25 Å, while QH2_move refers to those simulations where Q-N2 distance was over 25 Å. Distances were measured between Arg:CZ, Lys:NZ, Glu:CD, and Asp:CG atoms. The orange shaded plots and circles represent ion pairs which are closed when Q stays at site 2, while the cyan shaded plots and circles represent ion pairs closed when Q moves towards site 4.

Arg108^NDUFS7^, which was identified to be in contact with Q at both sites 2 and 4 (Fig. 3), makes an ion pair with Glu206^ND1^ when Q resides at site 2. However, when QH2 moves towards site 4, the ion pair breaks, coinciding with Arg108^NDUFS7^ making a strong interaction to the headgroup. Simultaneously, Glu206^ND1^ establishes a new ion pair with Arg199^ND1^. This agrees with the recent high resolution structural data on complex I which shows the Arg108^NDUFS7^ - Glu206^ND1^ ion pair distance to increase significantly between the turnover and native structures, equivalent to site 2 and site 4 Q binding, respectively (Parey et al., 2021). In addition, complex I structures from *Ovis aries* show the ion pair to be closed when the decylubiquinone is bound at site 1, and open when it is bound at site 4 (Kampjut & Sazanov, 2020).

Glu206^ND1^ also makes an ion pair with Arg297^ND1^ when Q is close to site 2, but when it diffuses towards site 4, Arg297^ND1^ displaces to form a relatively stable ion pair with Asp203^ND1^. While this interaction is also present when Q is close to site 2, the ion pair appears to be stabilized by QH2 moving to site 4 (56% vs. 92% occupancy). Interestingly, Asp203 has been proposed to be a key residue for redox coupled proton pumping based on both experiments and simulations (Nuber et al., 2021; Parey et al., 2021; Sharma et al., 2015; Yoga et al., 2019).

Another ion pair which appears to have a higher occupancy when Q is stable at site 2 is between Lys56^ND1^ and Asp115^NDUFS7^. These residues, which are known to be important for the activity of complex I (Garofano, Zwicker, Kerscher, Okun, & Brandt, 2003; Zickermann, Barquera, Wikström, & Finel, 1998), are positioned away from the Q binding sites. Despite this, the ion pair dissociation appears to coincide very well with the Q movement, suggesting that long range conformational changes may also be important for Q diffusion. Conversely, Arg27^ND1^ and Asp101^NDUFS7^ ion pair forms when Q diffuses to site 4. Interestingly, this ion pair is not observed in the PDB 6GCS with Q modeled at site 2, but is seen in the PDB 6RFR where Q is resolved at site 4. We also point out that mutation of Asp101^NDUFS7^ to an alanine residue leads to a drastic drop in activity (Yoga et al., 2019), highlighting the potential importance of Asp101-associated ion pair in Q dynamics.

Overall, here we have identified central elements in the form of charged residues that rearrange as Q moves in the Q tunnel. The open/closed dynamics of ion-pairs observed in our MD simulations is also in excellent agreement with existing structural and biochemical data. Additionally, the ion pairs identified to be linked with Q movement were further analyzed in the S3 simulations where Q movement was not seen (Fig. 2). Overall, the ion pairs in the S3 simulations match the states when Q was stable at site 2 (Table S2), even when QH2 is modelled. This further explains why QH2 does not diffuse from site 2 towards site 4 in S3 simulations.

## Discussion

Here, microseconds long MD simulations are performed on the *Y. lipolytica* complex I structures in which quinone molecules have been proposed to bind in the Q tunnel. Simulation data based on PDB 6GCS reveal Qox binds in a stable conformation at site 2, while QH2 tends to move - from this position towards site 4. For Q site nomenclature, see (Haapanen et al., 2019). This suggests that an oxidized Q (Qox) molecule at site 2 waits for electron transfer from N2 FeS cluster, whereas QH2, formed after redox-coupled proton transfer (Sharma et al., 2015), departs the site. Upon one electron transfer from N2 FeS cluster, semiquinone species may form. Our MD data show semiquinone molecules are also mobile in the Q tunnel, but it is the neutral semiquinone (SQ) species that diffuses maximally, from site 2 towards the entrance of the Q tunnel (sites 4/5). Anionic SQ on the other hand is more trapped within the Q tunnel, and would eventually convert to double reduced double protonated quinol (QH2) before exiting the site. Overall, our data suggests that Qox prefers to reside at sites 1 and 2, whereas reduced (and protonated) species such SQ and QH2 prefer to diffuse away towards entrance sites (4 and 5).

The behavior of the Q species is different in the simulations of higher resolution structure PDB 7O6Y simulations, even though Q has been predicted to bind in similar location in lower resolution structure (PDB 6GCS). Interestingly, Qox is found to be less stable at site 2 in simulations of high-resolution structure PDB 7O6Y, instead it moves closer to the N2 cluster to bind at site 1. This position, which has previously not been characterized in *Y. lipolytica* complex I structure, is important, as the proximity to the N2 cluster likely enhances the efficiency of electron transfer. In addition, QH2 in these simulations is extremely stable at the structural site 2 position, and does not travel to site 4 as in simulations based on PDB 6CGS. Similarly, the neutral SQ species diffuses away from site 2, but not as far as in 6GCS-based simulations. This is likely due to the difference in the position of the conserved and conformationally mobile β1-β2^NDUFS2^ loop, which appears to be in a conformation that allows access to site 1, but blocks access to site 4 in PDB 7O6Y, while in PDB 6GCS the opposite conformation is observed. Our data support an important role of β1-β2^NDUFS2^ loop, in particular His95, in coupling Q dynamics in the tunnel. In addition, we identify amino acid residues that are central for Q dynamics (Table S1).

We also note that the two structures of complex I obtained under turnover conditions (PDBs 6GCS and 7O6Y) have vastly different resolutions, 4.3 Å and 3.4 Å, respectively. This, along with the differently modeled conformations of amino acid residues in the vicinity of Q (and its position) are also likely the contributing factors for differing Q behavior observed in simulations of these complexes.

Q binding site 4, which is located at the interface of the ND1 and NDUFS7 subunits, is in close proximity to the E channel, an area of highly-conserved charged residues which leads to the membrane interior and ultimately the antiporter-like subunits (Baradaran, Berrisford, Minhas, & Sazanov, 2013). It has been suggested that this area is important for the coupling of redox reaction to proton pumping (Galemou Yoga et al., 2021; Gutiérrez-Fernández et al., 2020; Haapanen & Sharma, 2021; Kaila, 2018) and several different Q species have been modeled and simulated at this site with multiscale computational approaches (Haapanen et al., 2019; Haapanen & Sharma, 2017; Röpke et al., 2021). It is thus noteworthy that both QH2 and SQ are seen to move to this position, suggesting that the movement of reduced Q species towards site 4 may be a part of the proton pumping mechanism. Movement of QH2 is accompanied by changes in the structure of the protein surrounding the Q tunnel, in particular rearrangement of several charge-charged interactions involving conserved loops of ND1, PSST and 49 kD subunits. These data indicate that it is not only the redox state of Q which is important in Q binding and dynamics, but also the changes in the protein structure, in particular conserved ion pairs, that occur concurrently. This is also supported by the simulations in which ion-pair interactions do not reassemble, as a result of which, the quinone molecule remains immobile and stable at its original binding location, notably in S3 simulations with QH2 modelled (see Table S2). The changes in charge-charge interactions have also been observed in the recent high-resolution structures of complex I in native and turnover conditions (Parey et al., 2021), and have been suggested to be related to the proton pumping mechanism of complex I. Overall, the ion-pair rearrangements seen in our simulations, which drive Q dynamics in the Q tunnel, can be considered to be central component of the proton pumping mechanism of complex I.

Previous EPR studies on *E. coli* complex I revealed EPR signals of semiquinone species that have been suggested to be central to the proton pumping mechanism (Narayanan, Leung, Inaba, Elguindy, & Nakamaru-Ogiso, 2015). The EPR signal corresponding to the semiquinone species observed at ~35 Å from the N2 cluster is in close agreement with neutral SQ population observed in our MD simulations. Even though there are suggestions that SQ species are extremely short lived and not relevant for the redox-coupled proton pumping mechanism of complex I (Wright, Fedor, Hirst, & Roessler, 2020), it is possible that under certain conditions neutral SQ forms and escapes the binding sites near N2 (sites 1/2) to the entrance binding sites (sites 4/5). Due to the proximity of neutral SQ bound to the lipid bilayer, it may react with the oxygen solubilized in the membrane and lead to the formation of reactive oxygen species (ROS). Such an electron leak to oxygen would be minimized in case of anionic semiquinone (SQ-), which is better trapped in the Q tunnel of complex I.

## Materials and methods

All-atom molecular dynamics simulations were performed using three structures of complex I from *Yarrowia lipolytica* (PDBs 6GCS (Parey et al., 2018), 6RFR (Parey et al., 2019) and 7O6Y (Parey et al., 2021)). Small model systems were constructed with only subunits close to the Q binding tunnel included (ND3, ND1, NDUFS2, NDUFS3, NDUFS7, NDUFS8). Missing backbone atoms were modelled using Modeller software (Šali & Blundell, 1993) (ND3 residues 35 to 48 in PDB 6GCS; ND3 residues 45 to 59 and 114 to 119 in PDB 7O6Y) and missing sidechain atoms were added using VMD PSFGEN tool (Humphrey, Dalke, & Schulten, 1996). Note several sidechains in PDB 6GCS β1-β2^NDUFS2^ loop were modelled due to being unresolved in the structure. The protein was placed in a POPC lipid bilayer using CHARMM-GUI (Jo, Kim, Iyer, & Im, 2008), and TIP3P water was added along with 100 mM concentration of Na^+^/Cl^-^ ions. The head group of quinone molecule with nine isoprene units (Q9) was placed at site 2 in 6GCS- and 7O6Y-based setups. In 6GCS based setups, the Q9 headgroup was placed to overlap with the position of DBQ head group, coordinates of which are provided separately in (Parey et al., 2018). In the 7O6Y simulations, we placed the Q head group at the structurally resolved DBQ position. Similarly, a Q9 molecule was placed based on structurally resolved quinone binding site (site 4) in 6RFR-based setups. In all simulations, the Q9 tail was placed in the tunnel and allowed to relax with constrains on all other atoms. All components were treated with CHARMM force field (MacKerell Jr et al., 1998), (Klauda et al., 2010). The parameters of quinone and iron sulfur clusters were obtained from previous studies (Galassi & Arantes, 2015), (Chang & Kim, 2009). All amino acids were modeled in their standard protonation states; histidine residues were kept neutral with δ nitrogen protonated and all lysine, arginine, glutamic acid, and aspartic acid residues were charged, except for Asp67 and Glu69 of ND3 subunit to prevent unnatural hydration at the boundary of protein truncation. To relax the long Q9 tail and remove any steric clashes, a steepest descent energy minimization with NAMD was carried out, with all heavy protein atoms fixed. All subsequent simulations were performed with GROMACS software (Abraham et al., 2015). First the systems were minimized, followed by a 100 ps NVT simulation and 1 ns NPT simulation, all performed with constraints on heavy protein atoms. Next, the constraints were removed and a subsequent minimization and 100 ps NVT were performed, followed finally by a 10 ns NPT simulation. The production runs were then initiated using the Nosé-Hoover thermostat (Nosé, 1984), (Hoover, 1985) and Parrinello-Rahman barostat (Parrinello & Rahman, 1981), with LINCS algorithm (Hess, 2008) implemented and electrostatic interactions calculated by PME (Darden, York, & Pedersen, 1993). Production runs were extended to the microseconds timescale, and several simulations replicates were performed. All trajectory analysis was performed using Visual Molecular Dynamics (Humphrey et al., 1996). Table 1 shows a list of all simulations performed in this study and their lengths. Our smaller model systems are found to be stable despite system truncation, as shown by RMSD of protein with respect to time (see Fig. S4 and also ref. (Yoga, Parey, et al., 2020)). It is noteworthy that simulations on higher resolution structure show smaller RMSD values.

**Table 1:** List of molecular dynamics setups presented in this study.

| Structure used / setup name | Binding site where Q9 molecule modeled | Q state | Length of simulations |
| --- | --- | --- | --- |
| 6RFR / S1 | site 4 | Qox | 2045 ns<br>2022 ns<br>2042 ns |
|  |  | SQ- | 2018 ns<br>2029 ns<br>1308 ns |
|  |  | SQ | 2108 ns<br>2025 ns<br>1001 ns |
|  |  | QH2 | 2042 ns<br>2046 ns<br>2063 ns |
| 6GCS / S2 | site 2 | Qox | 2227 ns<br>2044 ns<br>2039 ns |
|  |  | SQ- | 2035 ns<br>2025 ns<br>2065 ns |
|  |  | SQ | 2060 ns<br>2088 ns<br>2065 ns |
|  |  | QH2 | 3034 ns<br>2097 ns<br>2931 ns |
| 7O6Y / S3 | site 2 | Qox | 1089 ns<br>1082 ns<br>1056 ns |
|  |  | SQ- | 1010 ns<br>968 ns<br>999 ns |
|  |  | SQ | 971 ns<br>985 ns<br>987 ns |
|  |  | QH2 | 1062 ns<br>1361 ns<br>1112 ns |

## Acknowledgements

VS is thankful to Academy of Finland, Sigrid Jusélius Foundation, Jane and Aatos Erkko Foundation, Magnus Ehrnrooth Foundation and University of Helsinki for financial support. Center for Scientific Computing (CSC) Finland is acknowledged for high performance computing time.

## Author contributions

JL performed simulations, analyzed data, drew figures, and wrote the manuscript. OH performed simulations and contributed to manuscript writing. VZ analyzed data and wrote the manuscript. VS designed the project, analyzed data, and wrote the manuscript.

## References

Abraham, M. J., Murtola, T., Schulz, R., Páll, S., Smith, J. C., Hess, B., & Lindahl, E. (2015). GROMACS: High performance molecular simulations through multi-level parallelism from laptops to supercomputers. SoftwareX, 1, 19–25.

Agip, A.-N. A., Blaza, J. N., Fedor, J. G., & Hirst, J. (2019). Mammalian Respiratory Complex I Through the Lens of Cryo-EM. Annual review of biophysics, 48, 165–184.

Angerer, H., Nasiri, H. R., Niedergesäß, V., Kerscher, S., Schwalbe, H., & Brandt, U. (2012). Tracing the tail of ubiquinone in mitochondrial complex I. Biochimica et Biophysica Acta (BBA)-Bioenergetics, 1817(10), 1776–1784.

Baradaran, R., Berrisford, J. M., Minhas, G. S., & Sazanov, L. A. (2013). Crystal structure of the entire respiratory complex I. Nature, 494(7438), 443–448.

Chang, C. H., & Kim, K. (2009). Density functional theory calculation of bonding and charge parameters for molecular dynamics studies on [FeFe] hydrogenases. Journal of chemical theory and computation, 5(4), 1137–1145.

Chung, I., Wright, J. J., Bridges, H. R., Ivanov, B. S., Biner, O., Pereira, C. S.,… Hirst, J. (2022). Cryo-EM structures define ubiquinone-10 binding to mitochondrial complex I and conformational transitions accompanying Q-site occupancy. Nature Communications, 13(1), 2758. doi:10.1038/s41467-022-30506-1

Darden, T., York, D., & Pedersen, L. (1993). Particle mesh Ewald: An N· log (N) method for Ewald sums in large systems. The Journal of chemical physics, 98(12), 10089–10092.

Djurabekova, A., Galemou Yoga, E., Nyman, A., Pirttikoski, A., Zickermann, V., Haapanen, O., & Sharma, V. (2022). Docking and molecular simulations reveal a quinone-binding site on the surface of respiratory complex I. FEBS Letters. doi:10.1002/1873-3468.14346

Fedor, J. G., Jones, A. J., Di Luca, A., Kaila, V. R., & Hirst, J. (2017). Correlating kinetic and structural data on ubiquinone binding and reduction by respiratory complex I. Proceedings of the National Academy of Sciences, 201714074.

Fendel, U., Tocilescu, M. A., Kerscher, S., & Brandt, U. (2008). Exploring the inhibitor binding pocket of respiratory complex I. Biochimica et Biophysica Acta (BBA)-Bioenergetics, 1777(7), 660–665.

Galassi, V. V., & Arantes, G. M. (2015). Partition, orientation and mobility of ubiquinones in a lipid bilayer. Biochimica et Biophysica Acta (BBA)-Bioenergetics, 1847(12), 1560–1573.

Galemou Yoga, E., Schiller, J., & Zickermann, V. (2021). Ubiquinone Binding and Reduction by Complex I—Open Questions and Mechanistic Implications. Frontiers in Chemistry, 9, 266.

Garofano, A., Zwicker, K., Kerscher, S., Okun, P., & Brandt, U. (2003). Two aspartic acid residues in the PSST-homologous NUKM subunit of complex I from Yarrowia lipolytica are essential for catalytic activity. Journal of Biological Chemistry, 278(43), 42435–42440.

Grba, D. N., & Hirst, J. (2020). Mitochondrial complex I structure reveals ordered water molecules for catalysis and proton translocation. Nature Structural & Molecular Biology, 27(10), 892–900.

Gu, J., Liu, T., Guo, R., Zhang, L., & Yang, M. (2022). The coupling mechanism of mammalian mitochondrial complex I. Nature Structural & Molecular Biology, 29(2), 172–182. doi:10.1038/s41594-022-00722-w

Gutiérrez-Fernández, J., Kaszuba, K., Minhas, G. S., Baradaran, R., Tambalo, M., Gallagher, D. T., & Sazanov, L. A. (2020). Key role of quinone in the mechanism of respiratory complex I. Nature Communications, 11(1), 1–17.

Haapanen, O., Djurabekova, A., & Sharma, V. (2019). Role of Second Quinone Binding Site in Proton Pumping by Respiratory Complex I. Frontiers in chemistry, 7, 221.

Haapanen, O., Reidelbach, M., & Sharma, V. (2020). Coupling of quinone dynamics to proton pumping in respiratory complex I. Biochimica et Biophysica Acta (BBA)-Bioenergetics, 148287.

Haapanen, O., & Sharma, V. (2017). Role of water and protein dynamics in proton pumping by respiratory complex I. Scientific Reports, 7(1), 7747. doi:10.1038/s41598-017-07930-1

Haapanen, O., & Sharma, V. (2021). Redox-and protonation-state driven substrate-protein dynamics in respiratory complex I. Current Opinion in Electrochemistry, 100741.

Hess, B. (2008). P-LINCS: A parallel linear constraint solver for molecular simulation. Journal of Chemical Theory and Computation, 4(1), 116–122.

Hoover, W. G. (1985). Canonical dynamics: equilibrium phase-space distributions. Physical review A, 31(3), 1695.

Humphrey, W., Dalke, A., & Schulten, K. (1996). VMD: visual molecular dynamics. Journal of molecular graphics, 14(1), 33–38.

Jo, S., Kim, T., Iyer, V. G., & Im, W. (2008). CHARMM-GUI: a web-based graphical user interface for CHARMM. Journal of computational chemistry, 29(11), 1859–1865.

Kaila, V. R. (2018). Long-range proton-coupled electron transfer in biological energy conversion: Towards mechanistic understanding of respiratory complex I. Journal of The Royal Society Interface, 15(141), 20170916.

Kampjut, D., & Sazanov, L. A. (2020). The coupling mechanism of mammalian respiratory complex I. Science, 370(6516).

Klauda, J. B., Venable, R. M., Freites, J. A., O’Connor, J. W., Tobias, D. J., Mondragon-Ramirez, C.,… Pastor, R. W. (2010). Update of the CHARMM all-atom additive force field for lipids: validation on six lipid types. The journal of physical chemistry B, 114(23), 7830–7843.

MacKerell Jr, A. D., Bashford, D., Bellott, M., Dunbrack Jr, R. L., Evanseck, J. D., Field, M. J.,… Ha, S. (1998). All-atom empirical potential for molecular modeling and dynamics studies of proteins. The journal of physical chemistry B, 102(18), 3586–3616.

Moser, C. C., Farid, T. A., Chobot, S. E., & Dutton, P. L. (2006). Electron tunneling chains of mitochondria. Biochimica et Biophysica Acta (BBA)-Bioenergetics, 1757(9), 1096–1109.

Narayanan, M., Leung, S. A., Inaba, Y., Elguindy, M. M., & Nakamaru-Ogiso, E. (2015). Semiquinone intermediates are involved in the energy coupling mechanism of E. coli complex I. Biochimica et Biophysica Acta (BBA)-Bioenergetics, 1847(8), 681–689.

Nosé, S. (1984). A unified formulation of the constant temperature molecular dynamics methods. The Journal of chemical physics, 81(1), 511–519.

Nuber, F., Mérono, L., Oppermann, S., Schimpf, J., Wohlwend, D., & Friedrich, T. (2021). A quinol anion as catalytic intermediate coupling proton translocation with electron transfer in E. coli respiratory complex I. Frontiers in Chemistry, 9, 291.

Parey, K., Brandt, U., Xie, H., Mills, D. J., Siegmund, K., Vonck, J.,… Zickermann, V. (2018). Cryo-EM structure of respiratory complex I at work. eLife, 7, e39213.

Parey, K., Haapanen, O., Sharma, V., Köfeler, H., Züllig, T., Prinz, S.,… Vonck, J. (2019). High-resolution cryo-EM structures of respiratory complex I: Mechanism, assembly, and disease. Science Advances, 5(12), eaax9484.

Parey, K., Lasham, J., Mills, D. J., Djurabekova, A., Haapanen, O., Yoga, E. G.,… Zickermann, V. (2021). High-resolution structure and dynamics of mitochondrial complex I—Insights into the proton pumping mechanism. Sci. Adv., 7(46), eabj3221. doi:doi:10.1126/sciadv.abj3221

Parrinello, M., & Rahman, A. (1981). Polymorphic transitions in single crystals: A new molecular dynamics method. Journal of Applied physics, 52(12), 7182–7190.

Pavelka, A. C., Benes, P., Strnad, O., Brezovsky, J., Kozlikova, B., Gora, A.,… Biedermannova, L. (2012). Damborsky J: CAVER 3. 0: A tool for analysis of transport pathways in dynamic protein structures. PLoS Comput Biol, 8, e1002708.

Röpke, M., Riepl, D., Saura, P., Di Luca, A., Mühlbauer, M. E., Jussupow, A.,… Kaila, V. R. I. (2021). Deactivation blocks proton pathways in the mitochondrial complex I. Proceedings of the National Academy of Sciences, 118(29), e2019498118. doi:10.1073/pnas.2019498118

Šali, A., & Blundell, T. L. (1993). Comparative protein modelling by satisfaction of spatial restraints. Journal of molecular biology, 234(3), 779–815.

Sazanov, L. A. (2015). A giant molecular proton pump: structure and mechanism of respiratory complex I. Nature Reviews Molecular Cell Biology, 16(6), 375–388.

Sharma, V., Belevich, G., Gamiz-Hernandez, A. P., Róg, T., Vattulainen, I., Verkhovskaya, M. L.,… Kaila, V. R. (2015). Redox-induced activation of the proton pump in the respiratory complex I. Proceedings of the National Academy of Sciences, 112(37), 11571–11576.

Teixeira, M. H., & Arantes, G. M. (2019). Balanced internal hydration discriminates substrate binding to respiratory complex I. Biochimica et Biophysica Acta (BBA)-Bioenergetics, 1860(7), 541–548.

Tocilescu, M. A., Fendel, U., Zwicker, K., Dröse, S., Kerscher, S., & Brandt, U. (2010). The role of a conserved tyrosine in the 49-kDa subunit of complex I for ubiquinone binding and reduction. Biochimica et Biophysica Acta (BBA)-Bioenergetics, 1797(6-7), 625–632.

Uno, S., Masuya, T., Zdorevskyi, O., Ikunishi, R., Shinzawa-Itoh, K., Lasham, J.,… Miyoshi, H. (2022). Diverse reaction behaviors of artificial ubiquinones in mitochondrial respiratory complex I. Journal of Biological Chemistry, 102075. doi:https://doi.org/10.1016/j.jbc.2022.102075

Warnau, J., Sharma, V., Gamiz-Hernandez, A. P., Di Luca, A., Haapanen, O., Vattulainen, I.,… Kaila, V. R. (2018). Redox-coupled quinone dynamics in the respiratory complex I. Proceedings of the National Academy of Sciences, 115(36), E8413–E8420.

Wikstrom, M., Sharma, V., Kaila, V. R., Hosler, J. P., & Hummer, G. (2015). New perspectives on proton pumping in cellular respiration. Chemical reviews, 115(5), 2196–2221.

Wright, J. J., Fedor, J. G., Hirst, J., & Roessler, M. M. (2020). Using a chimeric respiratory chain and EPR spectroscopy to determine the origin of semiquinone species previously assigned to mitochondrial complex I. BMC Biology, 18(1), 54. doi:10.1186/s12915-020-00768-6

Yoga, E. G., Angerer, H., Parey, K., & Zickermann, V. (2020). Respiratory complex I– Mechanistic insights and advances in structure determination. Biochimica et Biophysica Acta (BBA)-Bioenergetics, 1861(3), 148153.

Yoga, E. G., Haapanen, O., Wittig, I., Siegmund, K., Sharma, V., & Zickermann, V. (2019). Mutations in a conserved loop in the PSST subunit of respiratory complex I affect ubiquinone binding and dynamics. Biochimica et Biophysica Acta (BBA)-Bioenergetics, 1860(7), 573–581.

Yoga, E. G., Parey, K., Djurabekova, A., Haapanen, O., Siegmund, K., Zwicker, K.,… Angerer, H. (2020). Essential role of accessory subunit LYRM6 in the mechanism of mitochondrial complex I. Nature Communications, 11(1), 1–8.

Zickermann, V., Barquera, B., Wikström, M., & Finel, M. (1998). Analysis of the Pathogenic Human Mitochondrial Mutation ND1/3460, and Mutations of Strictly Conserved Residues in Its Vicinity, Using the Bacterium Paracoccus d enitrificans. Biochemistry, 37(34), 11792–11796.

