## Supplementary text for "Tunnel dynamics of quinone derivatives and its coupling to protein conformational rearrangements in respiratory complex I"

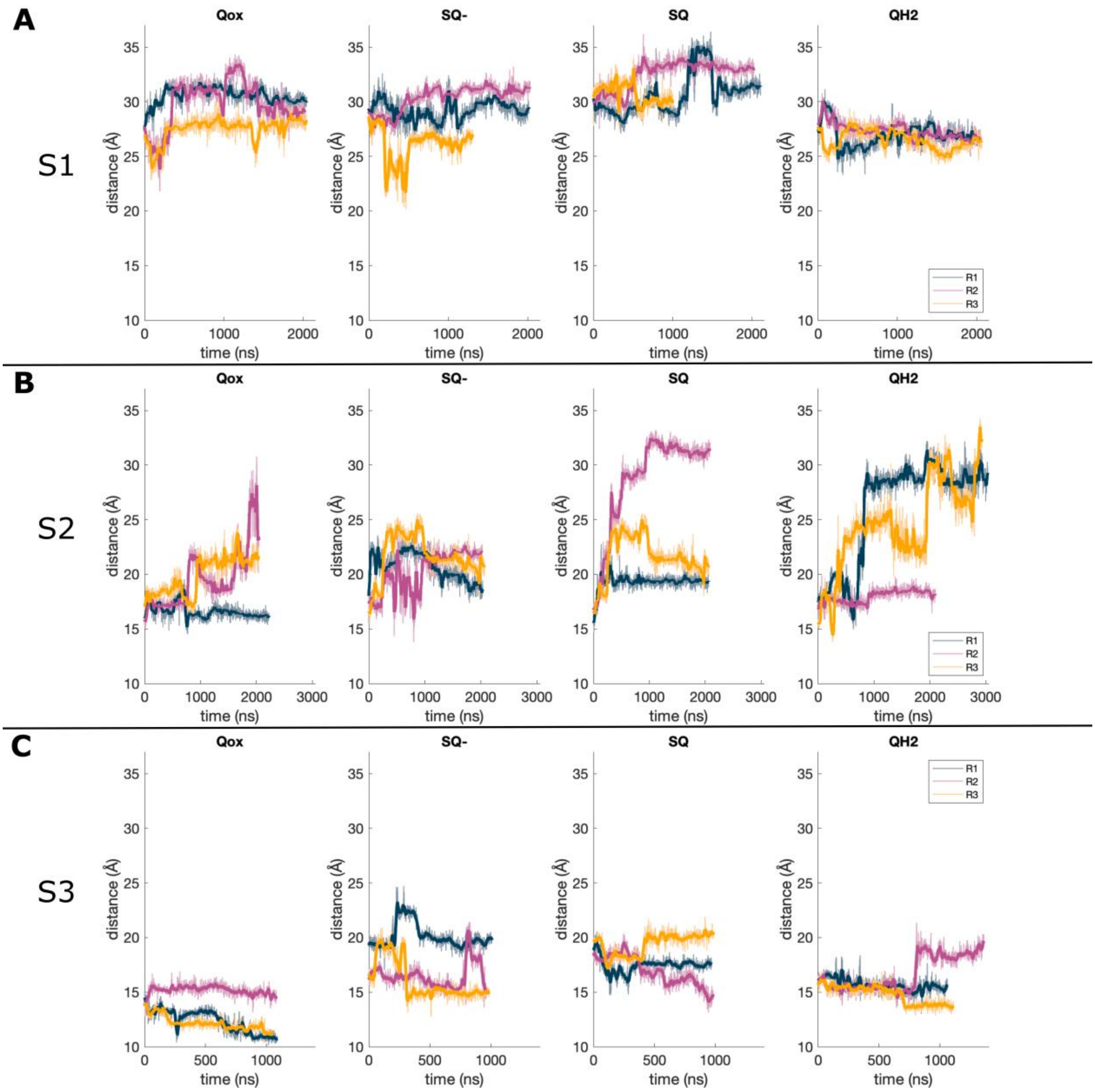

**Fig. S1**: Distance of Q9 head group (COM) from N2 cluster (COM) over time for S1 (A), S2 (B) and S3 (C) simulations. Different simulation replicas are shown in different colours, and the bold line shows a rolling average based on the previous 20 frames.

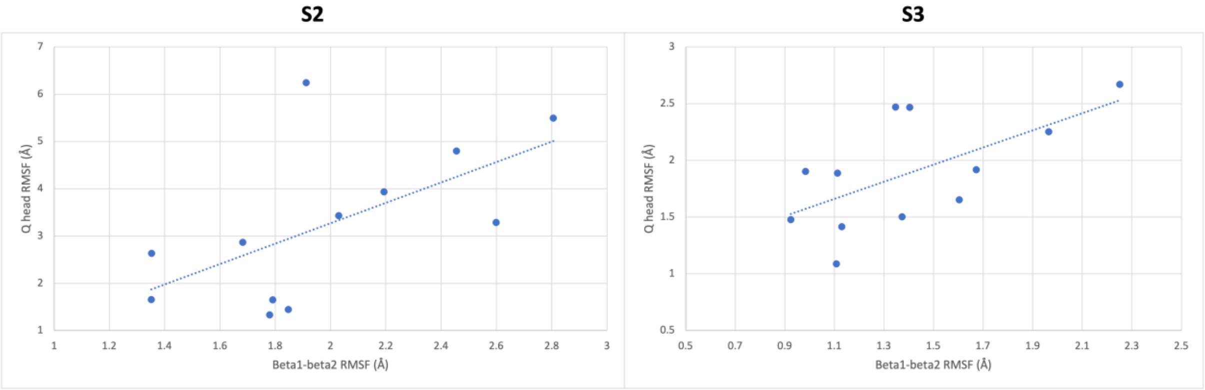

**Figure S2 –** Scatter plot of RMSF values of CA atoms of β1-β2$^{NDUFS2}$ loop and Q head atoms for S2 and S3 simulations.

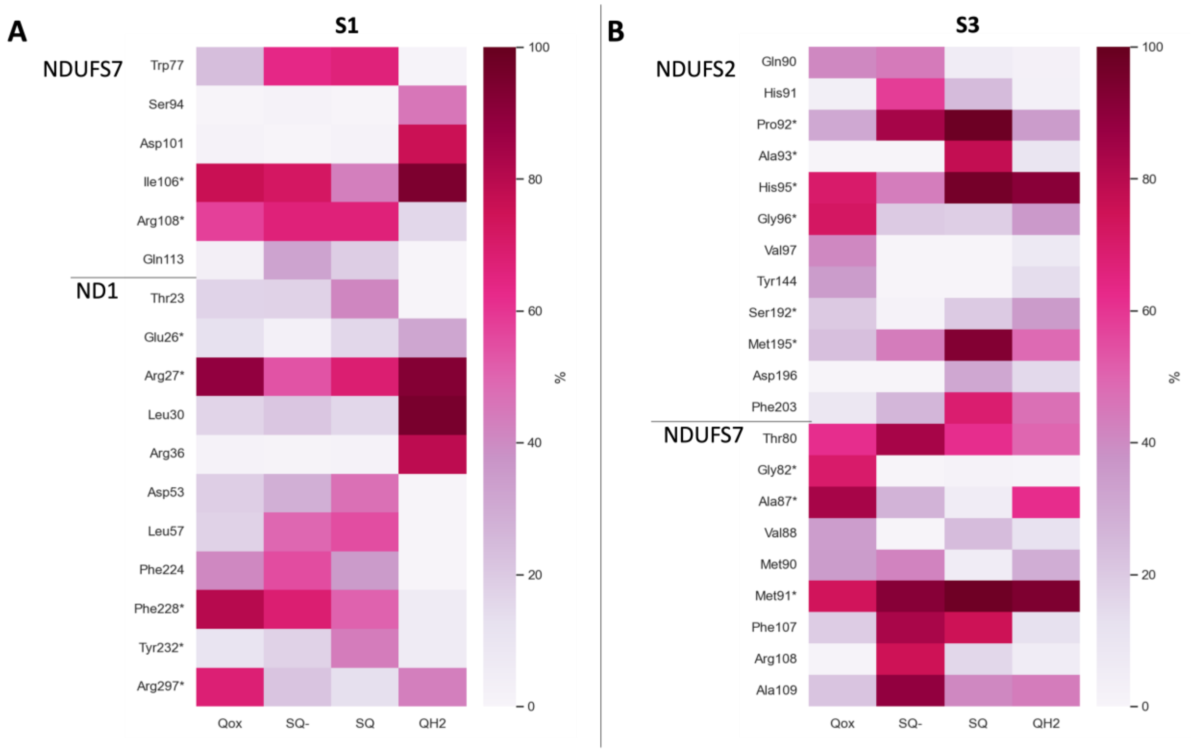

**Figure S3 –** Interactions between Q head group and protein residues in S1 simulations based on structure PDB 6RFR (A) and in S3 simulations based on high resolution structure PDB 7O6Y (B). The asterisk (*) denotes interactions between the Q head group and protein residues present in the respective structure.

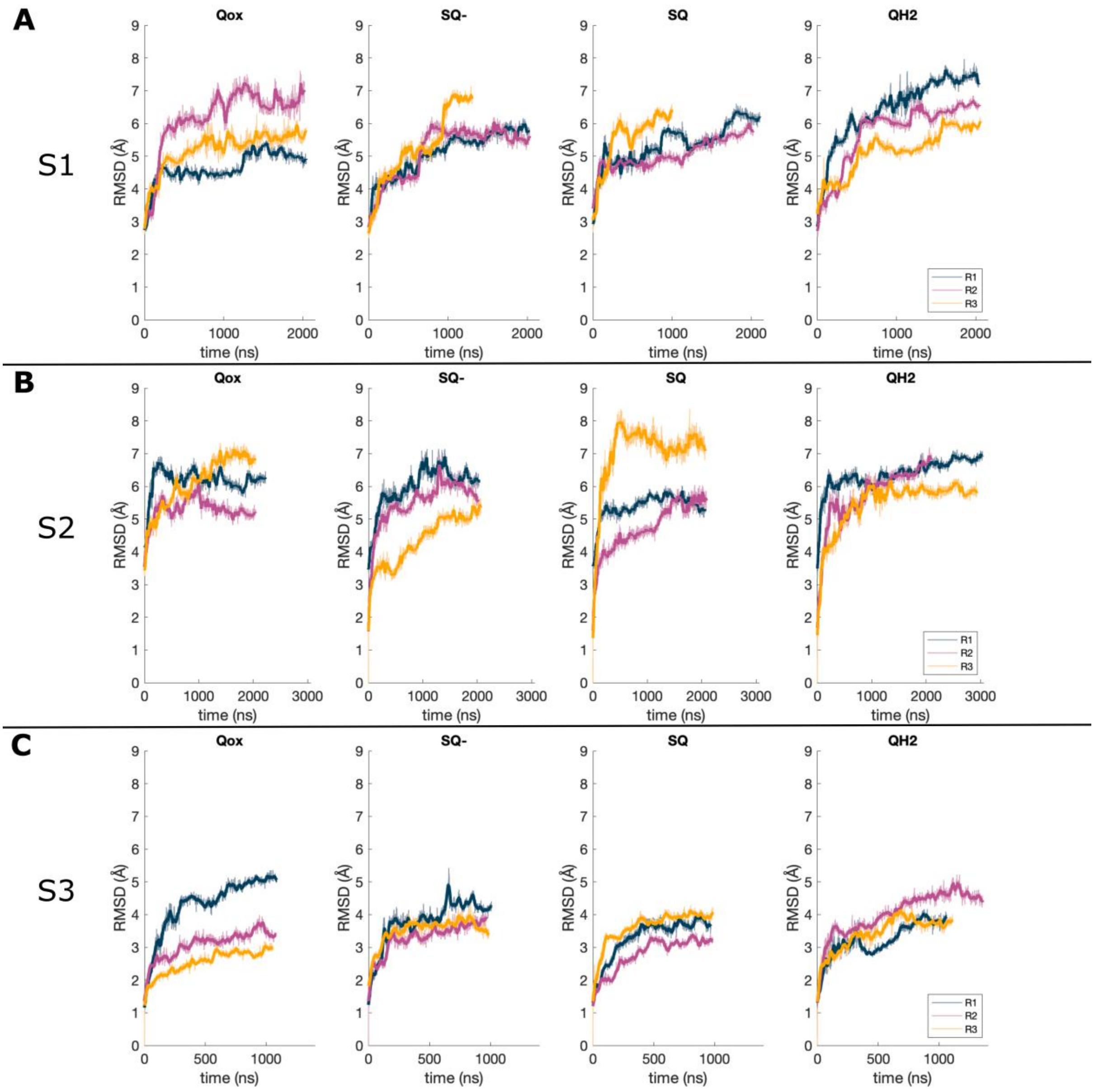

**Fig. S4**: RMSD of CA atoms over time for S1 (A), S2 (B) and S3 (C) simulations. Different simulation replicas are shown in different colours, and the bold line shows a rolling average based on the previous 20 frames.

| | | **Conservation** | **Mutation data** |
|---|---|---|---|
| **NDUFS2** | His91 | Conserved | H91A – reduced activity |
| | Pro92 | Conserved | |
| | Ala93 | Partly conserved | |
| | Ala94 | Partly conserved | A94I – decreased activity |
| | His95 | Conserved | H95A – decreased activity |
| | Gly96 | Conserved | |
| | Met195 | Partly conserved | M195F – decreased activity |
| | Leu200 | Partly conserved | |
| | Phe203 | Partly conserved | F203W – increased activity |
| | Leu204 | Partly conserved | |
| | Phe207 | Conserved | F207W – decreased activity |
| **NDUFS7** | Trp77 | Conserved | W77A/I/E – decreased activity |
| | Thr80 | Partly conserved | |
| | Ala87 | Partly conserved | |
| | Val88 | Partly conserved | V88L/M/F – decreased activity |
| | Met90 | Conserved | M90A/E – decreased activity |
| | Met91 | Partly conserved | M91C/K – decreased activity |
| | Ser94 | Partly conserved | S94A – no effect |
| | Asp101 | Partly conserved | D101A – decreased activity |
| | Ile105 | Partly conserved | I105A – no effect |
| | Ile106 | Partly conserved | I106F/A – decreased activity |
| | Phe107 | Partly conserved | F107A/P – decreased activity |
| | Arg108 | Conserved | R108A/E – decreased activity |
| | Ala109 | Partly conserved | A109G/S/C/L – decreased activity |
| | Asp115 | Conserved | D115N – decreased activity |
| **ND1** | Thr23 | Partly conserved | |

| | | | |
|---|---|---|---|
| | Glu26 | Conserved | E24K – human disease |
| | Arg27 | Conserved | R25Q – human disease |
| | Leu30 | Partly conserved | |
| | Arg36 | Conserved | R34H – human disease |
| | Asp53 | Conserved | D62E – *P. denitrifcans* - decreased activity |
| | Lys56 | Conserved | K67A - *P. denitrifcans* - decreased activity |
| | Leu57 | Partly conserved | |
| | Arg199 | Conserved | R195Q – human disease |
| | Asp203 | Conserved | D213A/E/N – E. *coli* – decreased activity |
| | Glu206 | Conserved | E216A – *E. coli* – decreased activity |
| | Phe224 | Partly conserved | |
| | Phe228 | Partly conserved | |
| | Tyr232 | Conserved | |
| | Ile294 | Partly conserved | |
| | Arg297 | Conserved | R291A – *E. coli* – decreased activity |

**Table S1 -** List of residues mentioned in the current work, along with their conservation and mutation data (if available). Mutation data for NDUFS2 is taken from (Angerer et al., 2012), for NDUFS7 from (Fendel, Tocilescu, Kerscher, & Brandt, 2008; Garofano, Zwicker, Kerscher, Okun, & Brandt, 2003; Yoga et al., 2019) and for ND1 from(Baradaran, Berrisford, Minhas, & Sazanov, 2013).

| | 6GCS | | 7O6Y | |
|---|---|---|---|---|
| **Ion pair** | **Qox (stay)** | **QH2 (move)** | **Qox** | **QH2** |
| R108 … E206 | Closed | Open | Open | Open |
| R199 … E206 | Mostly open | Mostly closed | Open | Mostly open |
| R297 … E206 | Mostly closed | Mostly open | Closed | Open |
| R297 … D203 | Mixed | Closed | Open | Open |
| R27 … D101 | Open | Closed | Mixed | Mixed |
| K56 … D115 | Mostly closed | Mostly open | Closed | Closed |

**Table S2** – Ion pairs identified to change with Q diffusion in the Q tunnel in S2 and S3 simulations. Data is shown for both Qox and QH2 species. See also main text Fig. 4. Closed was defined as the distance between Arg:CZ, Lys:NZ, Glu:CD, and Asp:CG atoms being < 5.5 Å.

### References

Angerer, H., Nasiri, H. R., Niedergesäß, V., Kerscher, S., Schwalbe, H., & Brandt, U. (2012). Tracing the tail of ubiquinone in mitochondrial complex I. *Biochimica et Biophysica Acta (BBA)-Bioenergetics, 1817*(10), 1776-1784.

Baradaran, R., Berrisford, J. M., Minhas, G. S., & Sazanov, L. A. (2013). Crystal structure of the entire respiratory complex I. *Nature, 494*(7438), 443-448.

Fendel, U., Tocilescu, M. A., Kerscher, S., & Brandt, U. (2008). Exploring the inhibitor binding pocket of respiratory complex I. *Biochimica et Biophysica Acta (BBA)-Bioenergetics, 1777*(7), 660-665.

Garofano, A., Zwicker, K., Kerscher, S., Okun, P., & Brandt, U. (2003). Two aspartic acid residues in the PSST-homologous NUKM subunit of complex I from Yarrowia lipolytica are essential for catalytic activity. *Journal of Biological Chemistry, 278*(43), 42435-42440.

Yoga, E. G., Haapanen, O., Wittig, I., Siegmund, K., Sharma, V., & Zickermann, V. (2019). Mutations in a conserved loop in the PSST subunit of respiratory complex I affect ubiquinone binding and dynamics. *Biochimica et Biophysica Acta (BBA)-Bioenergetics, 1860*(7), 573-581.
